# Automated high-throughput high-content autophagy and mitophagy analysis platform

**DOI:** 10.1101/412957

**Authors:** Jonathan Arias-Fuenzalida, Javier Jarazo, Jonas Walter, Gemma Gomez-Giro, Julia I. Forster, Paul M.A. Antony, Rejko Krueger, Jens C. Schwamborn

## Abstract

Autophagy and mitophagy play a central role in cellular homeostasis. In pathological conditions, the flow of autophagy and mitophagy can be affected at multiple and distinct steps of the pathways. Unfortunately, the level of detail of current state of the art analyses does not allow detection or dissection of pathway intermediates. Moreover, is conducted in low-throughput manner on bulk cell populations. Defining autophagy and mitophagy pathway intermediates in a high-throughput manner is technologically challenging, and has not been addressed so far. Here, we overcome those limitations and developed a novel high-throughput phenotyping platform with automated high-content image analysis to assess autophagy and mitophagy pathway intermediates.

## Introduction

Autophagy and mitophagy play central roles in normal development and disease^1,2^. Increasing interest and research in the field point out the need of pathway reconstitution tools and reliable quantification methods of autophagy and mitophagy^3,4^. Analyses of these processes have so far been conducted via low-throughput and semi-quantitative methods such as transmission electron microscopy, transient transfections, and western blot^5^. Current methodologies to assess autophagy and mitophagy impairments neglect the multiple structural intermediates of autophagy and mitophagy pathways. The implementation of technologies for identifying the successive developing structures in the autophagy and mitophagy pathway (staging) is necessary to dissect the impaired steps of the pathways. Such technologies enable to stratify and categorize pathologies that affect these crucial homeostasis pathways, and reveal new pharmaceutical targets^6^. The advent of genome editing tools has accelerated the development of genetically encoded reporters in human induced pluripotent stem (iPS) cells^7,8^. Additionally, the advancement of pH-responsive fluorescent proteins allows evaluating intracellular pH and interrogating specific subcellular compartments^9,10^. The combined use of genome editing tools and genetically encoded pH sensors enables the establishment of stable autophagy and mitophagy reporter lines. Automated high-throughput high-content imaging and analysis approaches have multiple advantages over conventional methods. Automated tools allow applying classification algorithms to specimens in an unbiased manner, and provide high statistical power. Importantly, those tools provide an insight to the flow of these pathways that would otherwise be missed in population-based analyses. Here, we developed and validated such automated tools to assess autophagy and mitophagy intermediates. Firstly, we combined TALEN technologies and the Rosella-reporter systems to engineer stable- tagged reporter-stem cells, advancing the detail of reporting. Secondly, this approach enabled us to acquire and quantify the respective pathway intermediates in our high-throughput phenotyping platform employing automated image analysis.

## Results

### Reconstruction of autophagy and mitophagy pathway intermediates

Making use of genetically encoded pH sensors and genome editing tools, we have engineered an iPSC line to monitor autophagy and mitophagy. The autophagy sensor Rosella-LC3 allows the identification of pre-autophagosomal structures such as phagophores, and transient structures such as isolation membranes and autophagosomes (Fig. 1A, C and E). Following autophagosome fusion with lysosomes, the internal membrane bound LC3 is degraded, giving rise to early and then late autolysosomes (Fig. 1A, C and E) that converge to lysosomes and re-enter the autophagy cycle^11,12^. The mitochondrial sensor ATP5C1-Rosella allows the quantification of the rate of mitophagy events (Fig. 1B, D, F) as accounted by acidic DsRed^pos^pHluorin^neg^ vesicles derived from degraded mitochondria. Using pattern recognition algorithms, we automatically identified and categorized the subcellular structures observed during autophagy (Fig. 1G). Autophagosomes and early autolysosomes are transient intermediates with low frequency levels, compared to the frequency of events at the extremes of the pathway: phagophores and late autolysosomes (Fig. 1G). Autophagic-vacuoles comprise all autophagosomes and autolysosomes in a cell^5^. The ratio between phagophores and autophagic-vacuoles accounts for the autophagy-rate-constant intrinsic to the cell line in basal conditions. In the here investigated example, the autophagy rate was close to 0.4 in a steady state.

**Fig 1:**
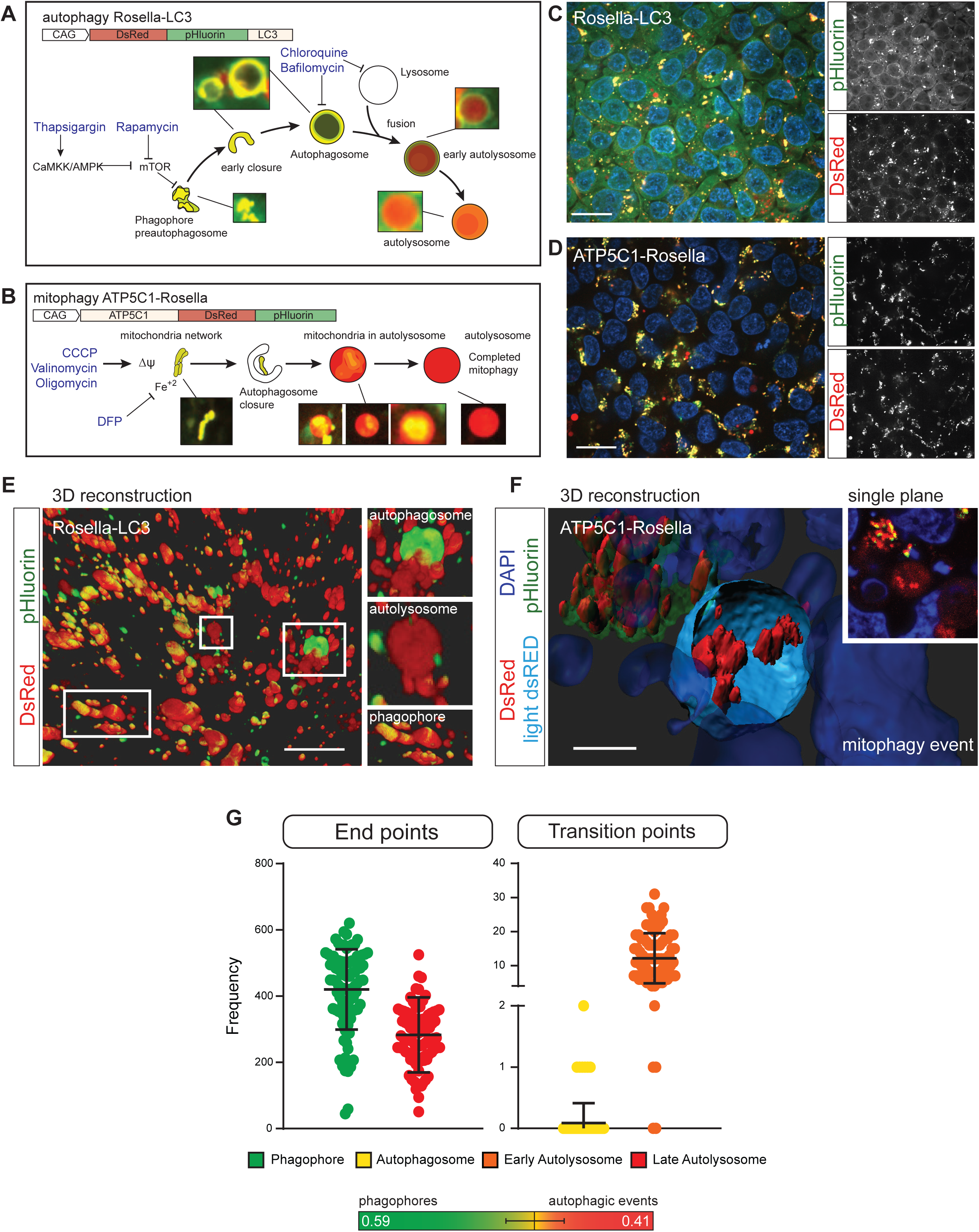
Genetically encoded Rosella-LC3 and ATP5C1-Rosella systems allow monitoring of the complete autophagy and mitophagy pathway. (**A**) Structure of the Rosella autophagy reporter system. It is possible to identify phagophores, autophagosomes, and autolysosomes. Small molecule modulators of autophagy can interrogate the autophagy responsiveness. (**B**) Structure of the Rosella mitophagy reporter system. It is possible to determine mitochondrial network structure and mitophagy events. Small molecule mitochondria stressors can test the mitophagy capacity. (**C**) Representative field for Rosella-LC3 line. The pHluorin and DsRed channels are shown separately. Scale bar, 10µm. (**D**) Representative field for ATP5C1-Rosella line. The pHluorin and DsRed channels are shown separately. Scale bar, 10 µm. (**E**) 3D reconstruction based on the Rosella-LC3 line. The insets show DsRed^pos^pHluorin^pos^ autophagosome structures, DsRed^pos^pHluorin^neg^ autolysosome structures, and DsRed^pos^pHluorin^pos^ phagophores. Scale bar, 10 µm. (**F**) 3D reconstruction based on the ATP5C1-Rosella line. An autolysosome structure with an ongoing mitophagy event is shown. The autolysosome appears with an equatorial cross section and the light DsRed volume is represented in cyan. The residual mitochondria inside the autolysosome are pHluorin^neg^, maintaining the pH-resistant DsRed signal. A phagophore DsRed^pos^pHluorin^pos^ cluster is located in the upper left-hand corner. In the inset, a single plane overlay is shown. The event was observed upon addition of 5 µM valinomycin. Scale bar, 4 µm. (**G**) Absolute quantification of phagophores, autophagosomes, early autolysosomes and late autolysosomes per field. At seeding density of as 600k cells/cm2, a range of 40-60 cells per field was obtained. Plus, quantification of the autophagy reaction rate constant.

### Autophagy image analysis decision tree

As shown in (Fig. 2), an automated computational image analysis workflow for the resulting multichannel 3D images was implemented. First, the raw images (Fig. 2A-C, pHluorinImRaw and dsRedImRaw) were flatfield corrected on the basis of reference images from an adjustment plate. The flatfield corrected images were deconvolved using the deconvblind function (Fig. 2D-E, pHluorinDeconvolved and dsRedDeconvolved). The number of iterations was set to 10 and the initial estimate of the point spread function was generated with the PSFGenerator tool^13^. In the PSFGenerator, the Richards & Wolf 3D optical model was used. The detailed parameter settings are summarized in Table S1.

**Fig 2:**
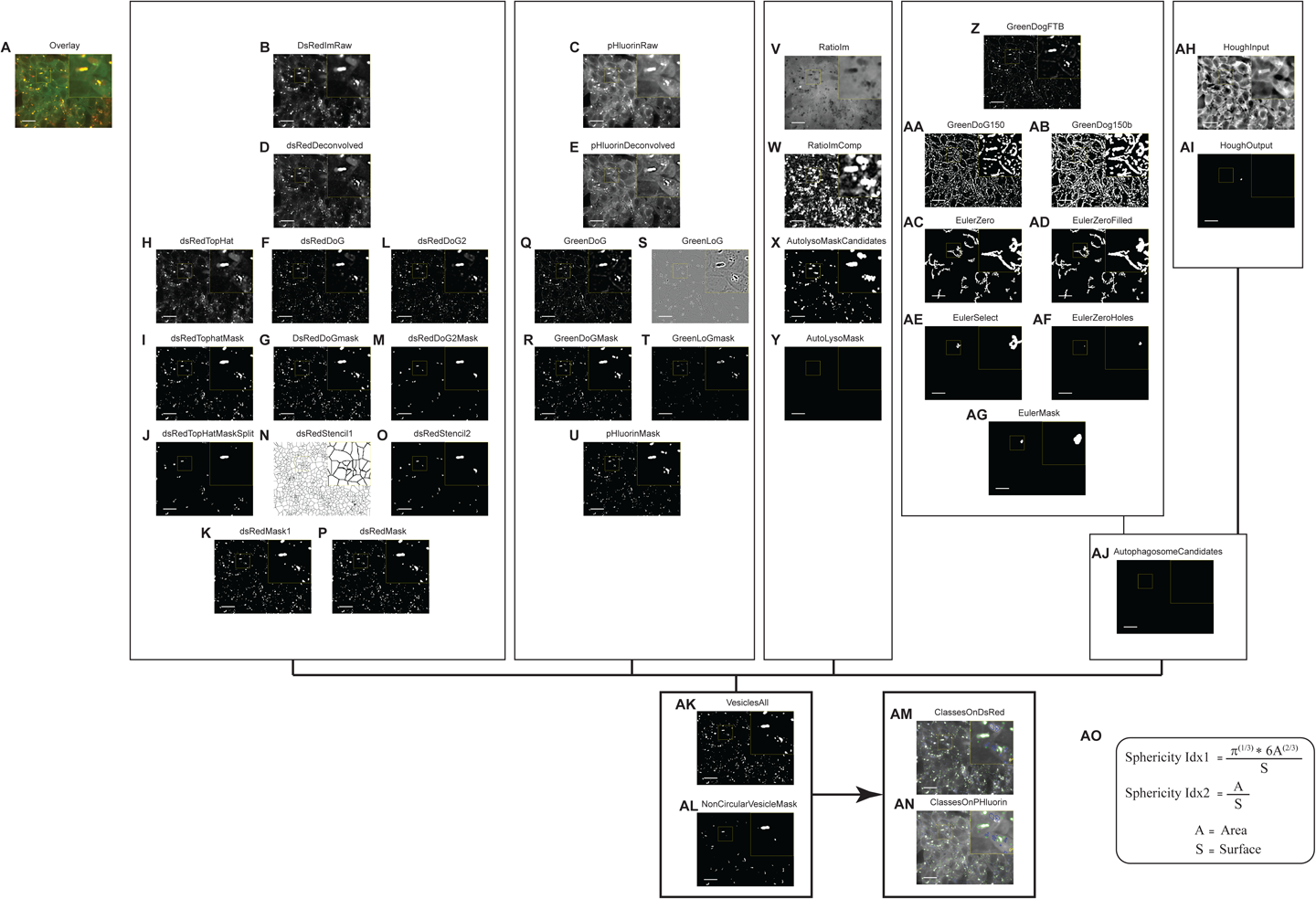
Image analysis workflow for autophagy Rosella-LC3 reporter line. (**A**) Overlay of raw DsRed and pHluorin channels. (**B**) Raw image for DsRed, dsRedImRaw. (**C**) Raw image for pHluorin, pHluorinImRaw channel. (**D**) dsRedDeconvolved. (**E**) pHluorinDeconvolved. (**F**) dsRedDoG. (**G**) dsRedDoGmask. (**H**) dsRedTopHat. (**I**) dsRedTopHatMask. (**J**) dsRedTopHatMaskSplit. (**K**) dsRedMask1. (**L**) dsRedDoG2. (**M**) dsRedDoG2Mask. (**N**) dsRedStencil1. (**O**) dsRedStencil2. (**P**) dsRedMask. (**Q**) GreenDoG. (**R**) GreenDoGMask. (**S**) GreenLoG. (**T**) GreenLoGmask. (**U**) pHluorinMask. (**V**) RatioIm. (**W**) RatioImComp. (**X**) AutoLysoMaskCandidates. (**Y**) AutoLysoMask. (**Z**) GreenDoGFTB. (**AA**) GreenDoG150. (**AB**) GreenDoG150b. (**AC**) EulerZero. (**AD**) EulerZeroFilled. (**AE**) EulerSelect. (**AF**) EulerZeroHoles. (**AG**) EulerMask. (**AH**) HoughInput. (**AI**) HoughOutput. (**AJ**) AutophagosomeCandidates. (**AK**) VesiclesAll. (**AL**) NonCircularVesicleMask. (**AM**) All vesicles mapped on DsRed channel. (**AN**) All vesicles mapped in pHluorin channel. Diameters shown in the results section correspond to the major axis length of the respective vesicles. Scale bars indicate 20 µm and 3x zoomed insets are highlighted with yellow boxes. **(AO)** Formula for calculating the sphericity indices used in the analysis.

After deconvolution of the DsRed channel, differences of Gaussians were computed in order to highlight DsRed^pos^ vesicles via spatial bandpass filtering. For the detection of small DsRed^pos^ vesicles, a foreground image was computed via convolution with a Gaussian filter of size and standard deviation (GF-S-SD) 20 pixel and 1 pixel, respectively. The subtracted background image was returned from convolution with GF-S-SD 20 pixel and 7 pixel (Fig. 2F, dsRedDoG). The mask of small DsRed^pos^ vesicles was defined via thresholding (>400) (Fig. 2G, dsRedDoGmask). To further improve the sensitivity of detection for DsRed^pos^ vesicles, this approach was complemented with a top-hat filtering approach. Top-hat filtering of dsRedDeconvolved was done using the imtophat function and a disk shaped structuring element of radius 25 pixel (Fig. 2H, dsRedTopHat). Thresholding (>1200) returned the corresponding mask (Fig. 2I, dsRedTopHatMask). To refine dsRedTopHatMask, connected components with more than 500 pixels, overlapping with more than 10% of its pixels with the dsRedDoGmask, were substituted with the corresponding pixels in dsRedDoGmask (Fig. 2J, dsRedTopHatMaskSplit).

Both, dsRedDoGmask and dsRedTopHatMaskSplit were combined using Boolean OR logic (Fig. 2K, dsRedMask1). To confirm the detection of DsRed^pos^ vesicles, a second difference of Gaussians was computed. Here the foreground image was convolved with a GF-S-SD of 11 pixel and 1 pixel while the background image was convolved with a GF-S-SD of 25 pixel and 6 pixel. The image resulting from subtraction of the background image from the foreground image (Fig. 2L, dsRedDoG2) was thresholded with gray tone value 1000 and objects were filtered by size between 200 and 2000 pixels (Fig. 2M, dsRedDoG2Mask). To split the vesicles, dsRedMask1 was used as a seed for a Euclidean distance transform and then the watershed function was applied. The resulting watershed mask (Fig. 2N, dsRedStencil1) was elementwise multiplied with the confirmative dsRedDoG2Mask (Fig. 2O, dsRedStencil2). The final DsRed^pos^ mask (Fig. 2P, dsRedMask) was computed via Boolean operation by pooling all pixels which were either present in dsRedMask1 or in dsRedStencil2.

To add more sensitivity for the detection of non-acidic vesicles, segmentation steps based on the deconvolved pHluorin channel were added. First, a difference of Gaussians was computed to segment larger vesicles. The GF-S-SD used to compute the foreground image was 100 pixel and 1 pixel, and the subtracted background image was convolved with a GF-S-SD of 100 pixel and 5 pixel (Fig. 2Q, GreenDoG). The mask of big green fluorescent vesicles was returned from intensity thresholding (>1000) (Fig. 2R, GreenDoGMask). To detect edges in the deconvolved pHluorin channel, a Laplacian of Gaussian filter with size 20 pixel and standard deviation 1 pixel was applied (Fig. 2S, GreenLoG), and pixels with values <-2000 were returned into the mask of edges (Fig. 2T, GreenLoGmask). The mask combining all vesicles detected in the pHluorin channel was computed via Boolean OR operation where the mask of big green fluorescent vesicles and the mask of edges were merged and connected components with less than 10 pixels were removed (Fig. 2U, pHluorinMask).

Ratio images (Fig. 2V, RatioIm) were computed by applying convolution GF-S-SD of 5 pixel and 2 pixel to the raw pHluorin and the raw DsRed channel. The ratio image was computed by elementwise division between the blurred pHluorin image and the blurred DsRed image.

To detect potentially missed autolysosomes, RatioIm was complemented using the imcomplement function (Fig. 2W, RatioImComp), and top-hat filtered using a disk-shaped structuring element of radius 15 pixel. Connected components above threshold 1.5 (Fig. 2X, AutoLysoMaskCandidates) were further validated as autolysomes by comparing their green fluorescence with the green fluorescence in the neighborhood defined by dilatation with a disk shaped structuring element of radius 7 pixel. Only when the neighborhood was at least 50% brighter than the vesicle and the vesicle volume was larger than 100 pixel, it was retained as an autolysosome candidate (Fig. 2Y, AutoLysoMask).

To detect autophagosomes based on their hollow vesicle property, an algorithm combining Fourier transform and Euler filtering was implemented. First, the GreenDoG image was plane by plane filtered with a Butterworth high pass filter in the Fourier transform domain. The cutoff frequency of the Butterworth filter was set to 10 and the order of the filter was set to 5. The resulting image (Fig. 2Z, GreenDoGFTB) was binarized via thresholding (>150) (Fig. 2AA, GreenDoG150). The resulting mask was maximum projected along the z axis. Objects with less than 20 pixel were removed, and median filtering using a 3 × 3 structuring element was applied. To prepare for the downstream analysis of connected components, the mask was further opened with a disk shaped structuring element of radius 1 pixel (Fig. 2AB, GreenDoG150b). Euler numbers were used to detect connected components containing holes. Indeed, an Euler number of 0 indicates that a connected component contains exactly one hole. Only connected components with Euler number 0 were retained (Fig. 2AC, EulerZero). For further filter refinement, the proportions between object and hole size were evaluated. For that purpose the imfill function was used in order to create a filled mask (Fig. 2AD, EulerZeroFilled). In the proportion filter, only objects with a ratio between area and filled area larger than 1.01, and a difference between filled area and area larger than 20 were retained (Fig. 2AE, EulerSelect). To segment the holes, the mask was inverted and the background was removed by applying a size threshold (10000 pixels) (Fig. 2AF, EulerZeroHoles). Next, the imreconstruct function was used, with the hole mask described above as the seed mask, and the FilledMask as the limiting mask. Finally, the shapes of autophagosome candidate vesicles were restored via morphological operations by applying image opening with a disk shaped structuring element of radius 5 pixel. For 3D reconstruction, only those planes which were already positive in GreenDoG150 were retained (Fig. 2AG, EulerMask).

To detect remaining autophagosomes, Hough transforms for circle detection were applied to the raw pHluorin channel. To minimize false positive detections, the already identified vesicle pixels were substituted by low pass filtered pixels. To highlight the not yet detected circular vesicles, graytone erosion with a disk shaped structuring element of radius 2 pixel was applied (Fig. 2AH, HoughInput). For the detection of circles, the function imfindcircles was used, where the radius was set to a range between 3-30 pixel. To remove false positives from the Hough transform algorithm, each candidate circle was further analyzed with respect to its value in the raw green and red channels. Only candidate circles with a large median absolute deviation among raw pHluorin pixels (>20), and a low 0.9th quantile of raw DsRed^pos^ pixels (<300) were retained (Fig. 2AI, HoughOutput).

The mask of all autophagosome candidates was computed by combining autophagosome candidates detected via the Fourrier-Euler algorithm or the Hough algorithm. To minimize the number of false positive autophagosomes, filters based on size, shape, and the ratio between vesicle border and vesicle center intensities were applied. The allowed sizes were set to the range between 50-10,000 pixel (Fig. 2AJ, AutophagosomeCandidates). For shape evaluation, the vesicle surface was defined by erosion with a sphere shaped structuring element of radius 1 pixel. Two sphericity indices were computed as shown in Fig. 2 (Fig. 2AO). Only vesicles with SphericityIdx1> 1 and SphericityIdx2> 1.5 were retained. For classification, all vesicles detected via the different approaches shown above were combined via Boolean operations. In order to maintain vesicle splitting, the perimeter of the autophagosome mask was excluded from the pooled vesicle mask. To exclude DsRed^neg^ vesicles from the downstream analysis, the mean intensity in the raw DsRed channel was measured for all connected components. Only vesicles with a mean intensity above 300 and not touching the border of the image were considered for the classification (Fig. 2AK, VesiclesAll). For the remaining vesicles, the eccentricity was computed and all connected components with eccentricity >0.9 were collected in the mask named NonCircularVesicleMask (Fig. 2AL).

The classification of the segmented vesicles was designed to classify four vesicle types, namely phagophores, autophagosomes, early autolysosomes, and late autolysosomes. For this purpose, a progressive exclusion algorithm was implemented. First, vesicles overlapping with the autophagosome mask and not overlapping with the NonCircularVesicleMask were classified as autophagosomes. Three condition sets were defined to classify the remaining vesicles as phagophores. Case1: vesicles with at least 25% overlap with both the pHluorinMask and the DsRedMask, a median ratio above 2 and without overlap with the AutoLysoMask. Case2: the 3rd quantile of the green channel was above 7500 and the 3rd quantile of the red channel above 4000. Case3: the green fluorescence in the center of the vesicle was at least 25% brighter than at the vesicle’s surface. For the remaining vesicles that were not classified as autophagosomes or phagophores, two conditions were defined for the vesicle labeling as late autolysosome. Case1: vesicles with at least 25% overlap with the DsRedMask and less than 10% overlap with the pHluorinMask or, alternatively, a median ratio below or equal to 2. Case2: the green fluorescence was lower in the vesicle center than at the surface. Remaining vesicles, not classified as autophagosomes, phagophores, or late autolysosomes were defined as early autolysosomes if they had at least 25% overlap with the DsRedMask and less or equal to 25% overlap with the pHluorinMask, or if they overlapped with the AutoLysoMask. Resulting segmented structures are shown for DsRed and pHluorin (Fig. 2AM-AN).

### Human iPS cells present active and dynamic autophagy and mitophagy

We observed remarkable activity of the autophagy pathway in human iPS cells (Video S1-4). Indeed, autophagy has previously been associated with the maintenance of pluripotency and resistance to senescence in stem cells and proved essential for pre-implantation development^1^. Furthermore, we observed autolysosome vesicles interacting with the surface of the mitochondrial network and abundant mitophagy events (Video S5-7). This is in agreement with reported mechanisms^14,15^ in this cell type^16,17^.

### Responsiveness to a panel of autophagy and mitophagy modulators

Our 3D analysis algorithm enables absolute quantification of single autophagy events of cells in monolayer cultures. In order to validate this approach with traditional immunostaining, we performed high-content quantification of LC3-positive structures (Fig. 3A and B). Each line in the heatmap (Fig. 1B) represents the averaged amount of vesicles observed of a well. The resolution of conventional LC3-antibody staining lacks the ability to resolve the stage-specific structures in the autophagy pathway. In contrast to LC3-antibody assays, the reporter lines used in the present work enable classification based on acidic content. LC3 antibody quantification, in contrast, is limited to total vesicle count, lacking pathway intermediates (Fig. 3B). Using chloroquine, a known inhibitor of the autophagy pathway flow at the autophagosome level^18^, an increase in LC3 positive vesicles was observed in both cases, however the cells presenting the Rosella construct depict that this rise is due to an increase in phagophores and autophagosomes. This demonstrates that the blockage induced by chloroquine generates an accumulation of vesicles at the early stages of autophagy. In order to validate our combined systems of stable reporters and new automated quantification tools, we evaluated the responsiveness to autophagic and mitophagic stress and modulation. We used validated small molecule perturbations for this approach. Upon addition of the H^+^-ATPase inhibitor bafilomycin, acidification of lysosomes was impaired. In agreement with the expected blockage of trafficking^5^, we observed an increased level of autophagosomes, and a decreased abundance in autophagic-vacuoles (Fig. 3C). Likewise, chloroquine led to the accumulation of autophagosomes and a decrease in autophagic-vacuoles (Fig. 3C).

**Fig 3:**
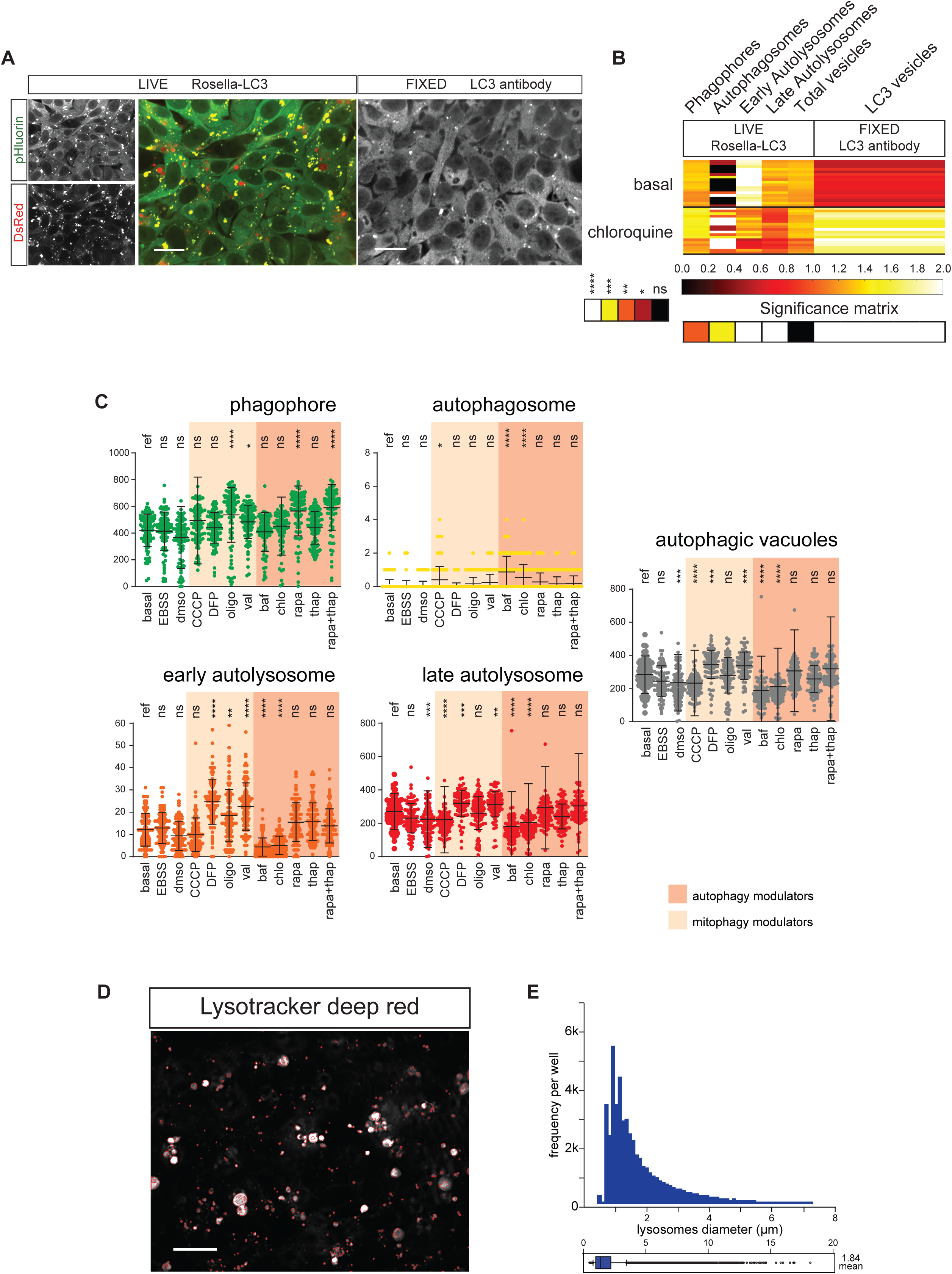
Quantification of autophagy structures before and after addition of mitophagy or autophagy modulators. (**A**) Representative fields of live Rosella-LC3 and fixed LC3 antibody. Scale bar, 20 µm. (**B**) Heatmap of scaled (0.0-2.0) category-mean normalized absolute frequency per well for autophagic structures for live Rosella and fixed LC3 antibody Rosella-LC3. Each line in the heatmap represents the averaged amount of vesicles observed of a well. Significance matrix of Kruskal-Wallis and Dunn’s multiple comparison test is shown below. Significance levels are *p<0.05, ** p<0.01, *** p<0.001, and **** p<0.0001. (**C**) Autophagic-vacuoles are quantified as the sum of autophagosomes, early autolysosomes, and late autolysosomes. Dunnett’s multiple comparison of means was performed for all conditions with respect to the basal reference (ref). Standard deviations are shown. Data represent three independent replicates. Significance levels are *p<0.05, **p<0.01, ***p<0.001 and ****p<0.0001. (**D**) Representative lysotracker staining. Lysosome mask on red perimeter. Scale bars indicate 20 µm. (**E**) Lysosome frequency and diameter in basal conditions.

Thapsigargin increases the intracellular concentration of Ca^2+^ and induces the activation of Calcium-Calmodulin-dependent-protein-kinase beta (CaMK-beta)^19^ which, through 5’AMP-activated-protein-kinase (AMPK), inhibits mechanistic-target-of-rapamycin (mTOR) and promotes autophagy^20^. Here, thapsigargin was used to quantify the CaMK-beta/AMPK dependent autophagy (Fig. 1A). However, no effect of treatment with Thapsigargin was observed (Fig. 3C).

It is known that mTOR complex 1 inhibits autophagy by its interaction with ULK1, resulting in reduced formation of the phagophore complex^20^. Rapamycin, an inhibitor of mTOR, allows quantification of the direct extent of autophagy controlled by this pathway. Oppositely to what we saw with the thapsigargin treatment, addition of rapamycin increased phagophore levels (Fig. 3C). This suggests that rapamycin modulates mTOR more efficiently compared to thapsigargin. We speculated that combined activation of CaMK-beta and repression of mTOR could have synergistic effects, however with the combined addition of thapsigargin and rapamycin we did not observe a further increase in phagophore formation (Fig. 3C). That is most likely due to the ubiquitous effects of the mTOR pathway^20^.

In order to evaluate the contribution of mitophagy to general autophagy, we used a panel of established mitophagy inducers^10^. Oligomycin, valinomycin, and CCCP are modulators of the mitochondrial membrane potential that induce mitochondrial stress and mitophagy. Only oligomycin and valinomycin induced an increase in phagophore levels (Fig. 3C). Stress induced by the high concentration of CCCP might have posed a toxic effect leading to the shut-down of the autophagy machinery, whereas a mild stress by DFP and valinomycin induced an increase in autophagic vacuoles (Fig. 3C). Known responses by mild stressors might be important for inducing an increase in Z’ values when doing assay development for drug screening purposes.

To determine the acidification capacity of autophagic-vacuoles, we complemented the autophagy and mitophagy reporters with a lysosomal dye (Fig. 3D and E) and developed a lysosome recognition algorithm (Fig. S1). This analysis would complete the phenotyping of the lysosmal-autophagosomal axis and allow discarding impairments in the flow of the autophagy pathway that are due to a decreased acidification capacity of autolysosomes.

### Assessment of mitochondria network volume and mitophagic vesicles volume

Next, we quantified the mitochondria network and mitophagic-vacuoles volume with the aid of pattern recognition algorithms (Fig. 4A-D). For the segmentation of mitochondria, the DsRed channel foreground signal was computed via convolution GF-S-SD of 50 pixel and 1 pixel (Fig. 4B). The subtracted background signal was computed via convolution with a GF-S-SD of 50 pixel and 2 pixel. Pixels with graytone values above 12 in this difference of Gaussians image (Fig. 4B, MitoDoG) were defined as mitochondrial pixels (Fig. 4B, MitoMask). MitophagyEvents were defined via a combination of green to red fluorescence ratio analysis and morphological filtering based on difference of Gaussians thresholding. First, 26 connected components within MitoMask were defined as mitophagy event markers (Fig. 4B, MitophagySeedMask) if the mean ratio value within the connected components was below 0.6. To refine the shape of the detected mitophagy events, the imreconstruct function was used with the parameters MitophagySeedMask and MitophagyLimitingMask (Fig. 4B). The MitophagyLimitingMask was defined by pixel values above 50 in a difference of Gaussians of the DsRed channel (Fig. 4B, MitophagyDoG). MitophagyDoG was defined by GF-S-SD of 50 pixel and 1 pixel for the foreground and GF-S-SD of 50 pixel and 5 pixel for the background. Resulting segmented mitochondria and mitophagic vacuoles are shown for DsRed and pHluorin. The disparity between the volumes of a mitochondrion and mitophagic vacuoles during the process of macroautophagy could be explained by different scenarios: several mitochondria prone to be degraded could be engulfed by the same vesicle, or addition of several lysosomes to the already formed autolysosome increases the vesicle size, or fusion of late autolysosome vesicles might occur (Fig. 4C). Upon addition of the mitochondrial stressor CCCP, the algorithm was able to detect an increase in the frequency of mitophagic events (Fig. 4D).

**Fig 4:**
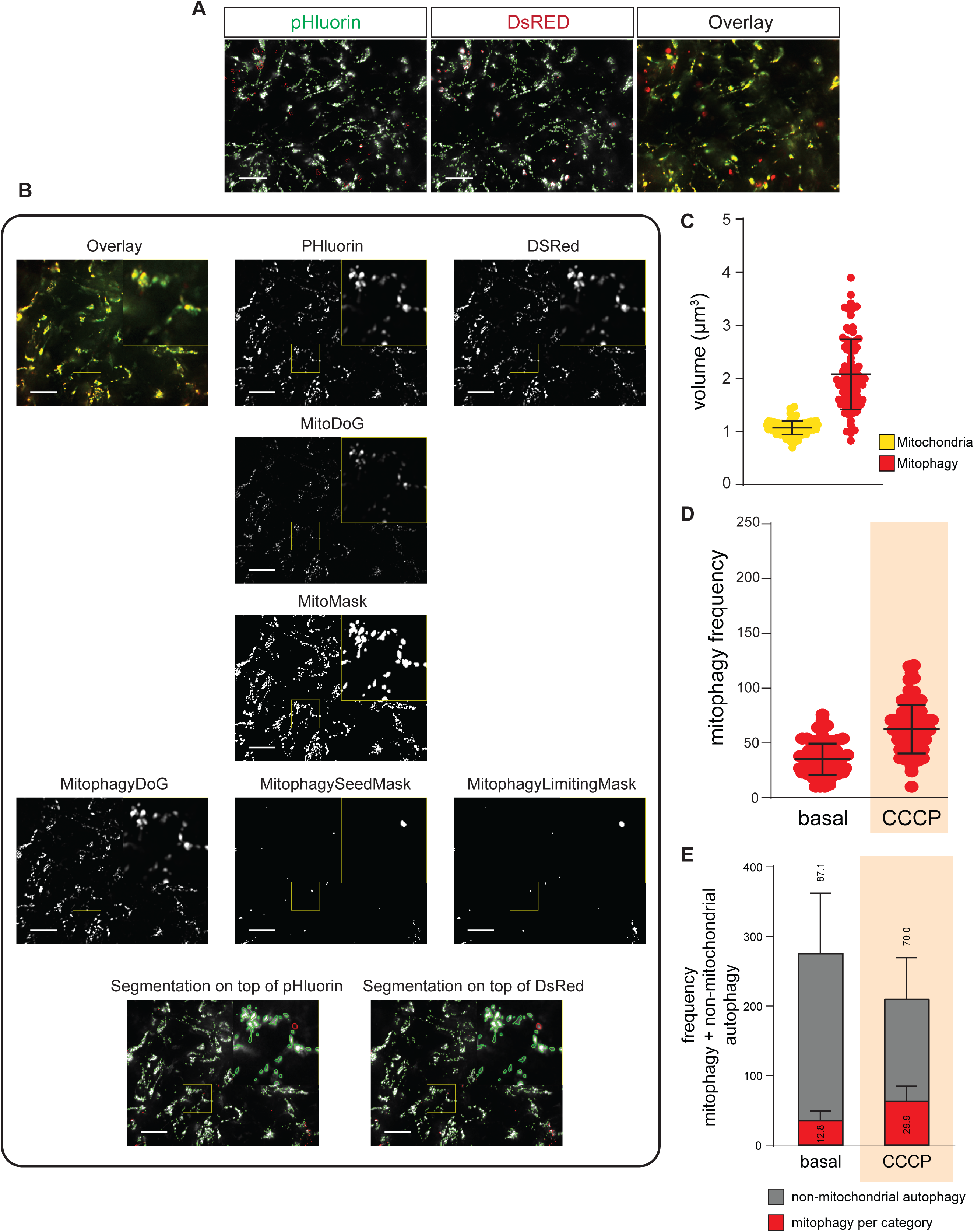
Evaluation of mitochondria network and mitophagic-vacuoles. (**A**) Representative images of ATP5C1-Rosella reporter. Mitochondria network mask on green perimeter and mitophagic-vacuole mask on red perimeter. Scale bars indicate 20 µm. (**B**) Image analysis workflow for mitophagy with ATP5C1-Rosella reporter line. Scale bars indicate 20 µm and 3x zoomed insets are highlighted with yellow boxes. (**C**) Mitochondrial and mitophagy volumes in basal conditions. (**D**) Mitophagy frequencies in basal condition and after mitochondrial stress induction. (**E**) Evaluation of mitophagy levels and distribution of autophagy resources upon mitochondrial stress. Percentages for mitophagy and non-mitochondrial autophagy are indicated.

Afterwards, we dissected the balance between mitophagy and general autophagy. The Rosella-LC3 sensor quantified the combined effect of non-mitochondrial autophagy and mitophagy. To model non-mitochondrial autophagy and to infer how phagocytic resources are distributed upon perturbations, we subtracted the ATP5C1- Rosella data from the Rosella-LC3 data. Then, we assessed the proportion of general autophagy and mitophagy resources in basal and stressed conditions. After CCCP treatment, the frequency of autophagic-vacuoles was reduced to 70% and mitophagy increased to 170% pointing that the autophagy machinery was shifted to mitochondria degradation (Fig. 4E).

## Discussion

The autophagy field faces the challenge of identifying the different stages of autophagy and mitophagy in a high-throughput manner^5^. The steps and sub-cellular structures of autophagy and mitophagy are important for the proper interpretation of phenotypic traits. This compartmentalization cannot be represented with bulk population analysis techniques such as western blotting. Here, we demonstrate a platform that accounts for vesicular compartmentalization using unbiased automated analysis in combination with automated high-content imaging, providing high statistical power and assay sensitivity. In the present work, we developed a high-throughput and high-content phenotyping platform for autophagy and mitophagy. This platform enables not only the evaluation of autophagy and mitophagy homeostasis in basal conditions but also for recognizing changes due to stress conditions, showing its potential application for detecting disease specific fingerprints. The platform presented was evaluated with an assortment of compounds to test the modulation of autophagy and mitophagy. The patterns observed are in agreement with the expected results of each modulator’s mechanism of action. We observed that lysosome fusion blockers bafilomycin and chloroquine increased the proportion of autophagosomes and reduced the proportion of autolysosomes as previously reported^5^. First, we validated our workflow by comparing it to immunostaining analysis of total LC3-positive structures. As mentioned above, the count of LC3- positive structures is lower than the total vesicle count quantified by the reporter. This can be explained by the degradation of LC3 epitope after the acidification of vesicles. On the contrary, in the reporter line the RFP signal remains visible enabling the quantification of late autophagy pathway structures. We show that the vesicles quantified by immunostaining correspond mainly to pre-acidification structures, principally phagophores (Fig. 1I). Comparing the LC3-positive structures from immunostaining and phagophores from the autophagy reporter line validates that both systems present the same autophagy signature upon chloroquine induction (Fig. 1I).

Our measurements were conducted on monolayer cultures with optimal seeding and treatment windows that maximized the autophagy and mitophagy pathways response. In this framework, the mild starvation modulator EBSS did not induce significant alteration of autophagy structure proportion (Fig. 3A). This demonstrates that the effects observed with the chemical modulators correspond to *bona fide* pathway responses. This approach offers a population-based analysis to determine the probabilistic flux level from basal conditions to treatment conditions by accounting for the different subcellular structures of the pathway. Due to the substantial level of automation, this platform can be used for genetic and chemical screening and thereby accelerate current endeavors in precision medicine.

## Materials and Methods

### Human iPS cell culture and electroporation

Human iPS cell line A13777 (Gibco) derived with non-integrative methods was used. Cells were maintained in Essential-8 media (Thermo Fisher cat no. A1517001) in feeder free culture condition on laminin 521 (BioLamina) or Matrigel (BD). Cell passage and dissociation was performed with accutase (Thermo Fisher cat no. A11105-01). Cells were electroporated with a Lonza 4D nucleofector system (Lonza V4XP-3024) according to the manufacturer’s instructions. After passage or electroporation, cells were cultured with 10µM Y27632 ROCK inhibitor (Sigma cat no. Y0503) for 24h.

### Autophagy and mitophagy reporter system

The pH sensor fluorescent protein pHluorin (F64L, S65T, V193G and H231Q) was fused to DsRed and the mitochondrial or autophagosomal targeting sequence ATP5C1 or LC3II as previously described^10^. The coding sequence was introduced into the AAVS1 safe harbor locus as previously described^8,21^ using the targeting donor (Addgene plasmid # 22075) and TALE nucleases (Addgene plasmid #35432 and #35431).

### Pathway contribution dissection

Reporter lines were treated with an assortment of compounds to dissect the stages of mitophagy and autophagy. In order to achieve a homogeneous monolayer of cells in each well, the optimal post-seeding time and cell density were determined for the lines used. The optimal density was identified as 600k/cm^2^ and the optimal post-seeding time as 8 hours, leaving a range of 40-60 cells per analyzed field. On 8 hours post-seeded cells, the minimal time required for assessing autophagy and mitophagy modulation were determined between 0.5 to 3 hours of treatment. The optimal imaging time of all compounds was identified as 3 hours after treatment, and was used for all experiments. Concentration gradients were tested to identify the minimal doses required to observe autophagy and mitophagy modulation without excessive cell toxicity after 3 hours of treatment, on 8 hours post-seeded cells. Final concentrations used from the ranges evaluated were 8 µM (8 µM-31.5 nM) bafilomycin A1 (Enzo); 8 µM (8 µM-31.5 nM) CCCP (Sigma cat no. C2759); 300 µM (300 µM-75 µM) chloroquine (Sigma cat no. C6628); 160 µM (160 µM-675 nM) DFP (Sigma cat no. D0879); 20 µM (20 µM-675 nM) oligomycin A (Sigma cat no. 75351); 160 µM (160 µM-675 nM) valinomycin (Sigma cat no. V3639); 40 µM (40 µM-156 nM) thapsigargin (Sigma cat no. T9033); and 160 µM (160 µM-675 nM) rapamycin (Sigma cat no. R8781). Minimal laser-exposure time was optimized for the samples in basal conditions.

### Immunostaining

Cells were fixed on 4% PFA in PBS and permeabilized with PBS triton-X 0.2%. Total human LC3 monoclonal antibody (MBL cat no. M152-3) was incubated at dilution 1:500 overnight. Secondary antibody was goat anti-rabbit alexa fluor 647 (Thermo cat no. A32733) and used at dilution 1:1000.

### Lysosome quantification and nuclear contrast

Cells under basal conditions were treated with deep red lysotracker (Thermo cat no. L12492) at a dilution of 1:1000 for 30 minutes. For nuclear staining, cells were treated with 20µM Hoechst 33342 for 10 minutes.

### Time-lapse live cell imaging

Culture dynamics and time lapse imaging was evaluated in a spinning disk CSU-X1 system (Zeiss) under controlled atmosphere conditions. Time-lapse imaging was performed for a single confocal plane. For three-dimensional pathway reconstruction, a single time point was evaluated. Reconstruction of 3D structures was performed with an Imaris (Bitplane) image processing 7.0 system.

### Microscopy for Rosella-LC3 and ATP5C1-Rosella

Confocal images were acquired on an Opera QEHS spinning disk microscope (Perkin Elmer) using a 60x water immersion objective (NA = 1.2). DsRed and pHluorin images were acquired in parallel using two cameras and binning 2. pHluorin was excited with a 488 nm laser and DsRed with a 561 nm laser. A 568 dichroic mirror was used to deviate the emitted light towards the corresponding cameras. pHluorin was detected on camera 1 behind a 520/35 bandpass filter and DsRed on camera 2 behind a 600/40 bandpass filter. For Rosella-LC3, five planes were set with 400 nm z-steps. For ATP5C1-Rosella, eleven planes were set with 400 nm z-steps. Scale of 1 pixel corresponds to 0.2152µm in all the cases described.

### Microscopy for the Lysotracker assay

Images were acquired on an Opera QEHS High content screening microscope using a 60x water immersion objective (NA = 1.2). Lysotracker deep red was excited with a 640 nm laser and detected with a 690/70 bandpass filter using camera binning 2. Z-stacks were defined to contain 11 planes with 400 nm z-steps.

### Image analysis for the Lysotracker assay

Deconvolution of raw images (Fig. S1A, LysoTDR) was done as described above according to the settings shown in table 1 (Fig. S1B, LysoTDR_deconvolved). For the segmented lysotracker positive vesicles, an algorithm with different morphological filters was implemented. First, a difference of Gaussians was computed using a GF- S-SD of 100 pixel and 1 pixel for convolving the foreground image and a GF-S-SD of 100 pixel and 5 pixel for convolving the subtracted background image (Fig. S1C, LysoTDR_DoG). A first mask with Lysotracker pixels was returned by thresholding (>2000) LysoTDR_DoG (Fig. S1E, LysoTracker_DoG_Mask). An additional detection option was implemented using convolution of LysoTDR_deconvolved with a Laplacian of Gaussian of GF-S-SD 20 pixel and 1 pixel (Fig. S1D, LysoTDR_LoG). The second mask of lysotracker pixels was returned by retaining pixels with graytone values <-2000 in LysoTDR_LoG (Fig. S1F, LysoTDR_LoG_Mask). The final mask of lysotracker stained vesicles was computed via Boolean OR combination of both masks and size exclusion of connected components with less than 10 pixels (Fig. S1G, LysoTDR_Mask). To compute the major axis length of each vesicle, LysoTDR_Mask was maximum projected and subsequently the function regionprops was used to extract the major axis length of each connected component in the projected mask.

## Acknowledgements

We thank Prof. J. Hejna and E. Berger for critical comments on the manuscript. We would like to thank Prof. T. Graham and A. Sargsyan from the University of Utah for kindly providing us with pHluorin constructs, Prof. R. Jaenisch from the MIT Whitehead Institute for providing the AAVS1 targeting vector, and Prof. F. Zhang from the McGovern Institute for Brain Research for providing the TALEN vectors. This project was supported by the LCSB pluripotent stem cell core facility. This project was funded by the Fonds National de la Recherche (FNR) Luxembourg (CORE, C13/BM/5791363). This is an EU Joint Program - Neurodegenerative Disease Research (JPND) project (INTER/JPND/14/02; INTER/JPND/15/11092422). J.W., J.J. and J.F. were supported by FNR Aides à la Formation-Recherche (AFR). G.GG. was funded by the NCL-Stiftung (Hamburg, Germany).

## Author Contributions

J.AF. and G.GG. performed cloning. J.AF., J.W. and G.GG. performed knock-ins. J.AF., J.W., J.J. and P.A. performed automated image acquisition experiments. J.AF., J.J. and J.W performed compound library experiments. P.A. led automated image processing and pattern recognition programming. J.AF., J.J., J.W. and J.F. contributed to automatic image processing coding. P.A., J.AF., J.J. and J.W. performed analysis of results, wrote the manuscript, and organized the figures. J.S. supervised. All authors reviewed and agreed to the final version of the manuscript.

## Competing Interests

The authors declare no competing interest

